# Development of a Novel Bruton’s Tyrosine Kinase Inhibitor that exerts Anti-Cancer Activities Potentiates Response of Chemotherapeutic Agents In Multiple Myeloma Stem Cell-Like Cells

**DOI:** 10.1101/2022.03.10.483708

**Authors:** Weam Othman Elbezanti, Omar S. Al-Odat, Robert Chitren, Jaikee Kumar Singh, Sandeep Kumar Srivastava, Krishne Gowda, Shantu Amin, Gavin P. Robertson, Subash C. Jonnalagadda, Tulin Budak-Alpdogan, Manoj K. Pandey

**Affiliations:** Department of Biomedical Sciences, Cooper Medical School of Rowan University, Camden, NJ 08103, USA; Department of Biosciences, Manipal University Jaipur, Jaipur 303007, India; Department of Pharmacology, Penn State Hershey Cancer Institute, Penn State College of Medicine, 500 University Drive, Hershey, PA 17033, USA; Department of Chemistry and Biochemistry, College of Science and Mathematics, Rowan University, Glassboro, NJ 08028, USA; Department of Hematology, MD Anderson Cancer Center at Cooper, Cooper Health University, Camden, NJ 08103, USA

**Keywords:** Bruton’s tyrosine kinase, BTK inhibitor, multiple myeloma, hematological malignancies, multiple myeloma stem cell-like cells, acquired mutation

## Abstract

Despite recent improvements in multiple myeloma (MM) treatment, MM remains an incurable disease and most patients experience a relapse. The major reason for myeloma recurrence is the persistent stem cell-like population. It has been demonstrated that overexpression of Bruton’s tyrosine kinase (BTK) in MM stem cell-like cells is correlated with drug resistance and poor prognosis. We have developed a novel small BTK inhibitor, KS151, which is unique compared to other BTK inhibitors. Unlike ibrutinib, and the other BTK inhibitors such as acalabrutinib, orelabrutinib, and zanubrutinib that covalently bind to the C481 residue in the BTK kinase domain, KS151 can inhibit BTK activities without binding to C481. This feature of KS151 is important because C481 becomes mutated in many patients and causes drug resistance. We demonstrated that KS151 inhibits *in vitro* BTK kinase activities and is more potent than ibrutinib. Furthermore, by performing a semi-quantitative, sandwich-based array for 71-tyrosine kinase phosphorylation, we found that KS151 specifically inhibits BTK. Our western blotting data showed that KS151 inhibits BTK signaling pathways and is effective against bortezomib-resistant cells as well as MM stem cell-like cells. Moreover, KS151 potentiates the apoptotic response of bortezomib, lenalidomide, and panobinostat in both MM and stem cell-like cells. Interestingly, KS151 inhibits stemness markers and is efficient in inhibiting Nanog and Gli1 stemness markers even when MM cells were co-cultured with bone marrow stromal cells (BMSCs). Overall, our results show that we have developed a novel BTK inhibitor effective against the stem cell-like population, and potentiates the response of chemotherapeutic agents.

## INTRODUCTION

Multiple myeloma (MM) is a type of hematologic malignancy that accounts for 10% of all blood cancers and 1% of all cancers [1]. It is a clinically and biologically heterogeneous neoplasm and characterized by the uncontrolled expansion of malignant plasma cells (PCs) in the bone marrow (BM) [2]. Even though there is significant progress in MM patient survival in recent years with new treatments, almost all patients experience a relapse after their remission [3, 4].

MM stem cell-like cells are one of the key players in acquired drug resistance and are critical to relapse. Therefore, it is reasonable to believe that any effective treatment for MM will have to be effective at targeting and eliminating the pool of stem cell-like cells [5, 6]. Nonetheless, the cellular and molecular profile of MM stem cell-like cells is still controversial [7, 8]. It has been demonstrated that clonotypic CD138^neg^ plasma cells display self-renewal, tumor-initiating potential, and drug resistance, suggesting that they have stem cell-like properties [9]. In addition, it is possible that MM represents a number of biologically distinct diseases, each containing different stem cell populations [7]. For example, side population (SP) cells are accepted to have characteristics of stem cell-like cells [10, 11]. It was also demonstrated that CD19+CD27+CD138^neg^ with a memory B cell phenotype could engraft NOD/SCID mice during both primary and secondary transplantation [5]. Furthermore, both the SP and aldefluor assays were able to identify CD19+CD27+CD138^neg^ B cells within the peripheral blood of patients with MM. side population (SP). Even though there are many different types of stem cells in MM, CD138^neg^ or side population (SP) cells with high aldehyde dehydrogenase (ALDH) activity are thought to be the most representative of MM stem cells [9, 12–14]. In addition to CD138^neg^, Nanog, Myc, OCT4, and GLI1 are all-present on CD138^neg^ cells and can be used as additional markers of stemness [15, 16].

One of the critical pathways that has been identified as a potential target for MM stem cell-like cells is Bruton’s Tyrosine Kinase (BTK), a non-receptor tyrosine kinase [17, 18]. BTK plays an essential role in the differentiation and the development of B-lymphocytes. It is composed of five domains:1) the amino terminal pleckstrin homology (PH) domain, 2) the proline-rich Tec homology (TH) domain, the Src domains 3) SH3, 4) SH2 domain, and 5) the kinase domain (SH1) which is responsible for the catalytic activity [19] (**Figure 1A**). BTK contains two essential phosphorylation sites that are located in the SH3 domain (Y223) and in the kinase domain (Y551). BTK is activated through its phosphorylation at Y551 by other kinases such as Blk, Lyn, and Fyn followed by autophosphorylation at Y223 [20–22]. Following its activation, BTK phosphorylates phospholipase-Cγ2 (PLCγ2), and therefore induces Ca^2+^ mobilization, NF-κB, Akt, and MAP kinase pathway activation [23, 24]. BTK regulates several critical signal pathways that includes phosphatidylinositol 3-kinase (PI3K), and protein kinase C (PKC) [25]. This in turn regulates survival, proliferation, differentiation, and apoptosis [4]. Upregulation of BTK has been associated with increased metastasis, migration, poor prognosis, and relapse in malignancies [17, 26–28]. Recent studies have demonstrated that BTK is significantly expressed in MM primary cells, as well as its microenvironmental cells such as stromal, osteoclasts, and MM stem cell-like cells [17, 29]. BTK offers a strategy to overcome existing challenges in MM treatment. For example, BTK is upregulated in MM patients following relapse to standard-of-care chemotherapy bortezomib [30]. Bortezomib is a proteasomal inhibitor used as a frontline treatment for patients with MM but despite a positive initial response, all patients relapse [31, 32]. The potential mechanisms of resistance include upregulation of survival pathways, among them the activation of NF-κB and AKT [30, 33]. Additional work has demonstrated that the BTK inhibitor ibrutinib can inhibit NF-κB and that this effect is responsible for ibrutinib enhancing the cytotoxicity of bortezomib in MM models [34]. Ibrutinib, a first in class, once-daily oral inhibitor of BTK, and is approved by the FDA for the treatment of B cell malignancies [35, 36]. In patients with relapsed/refractory (R/R) MM, ibrutinib demonstrated encouraging activity in combination with dexamethasone, with carfilzomib plus dexamethasone, and with bortezomib and dexamethasone [37–40]. Nonetheless, ibrutinib seems to be ineffective in the long term because of acquired mutations that often involve the mutations of Cys481 to Ser (C481S) in the kinase domain [41–44]. The C481S mutation disrupts the covalent binding between BTK and ibrutinib, which leads to a loss of inhibition of BTK enzymatic activity that ultimately results in ibrutinib resistance [41–46]. Though this mutation has not yet been reported in MM, likely due to sparse use of ibrutinib in this patient population, the correct BTK-targeting strategy needs to account for this possibility and an agent that can overcome the C481S mutation must be used.

**Figure 1:**
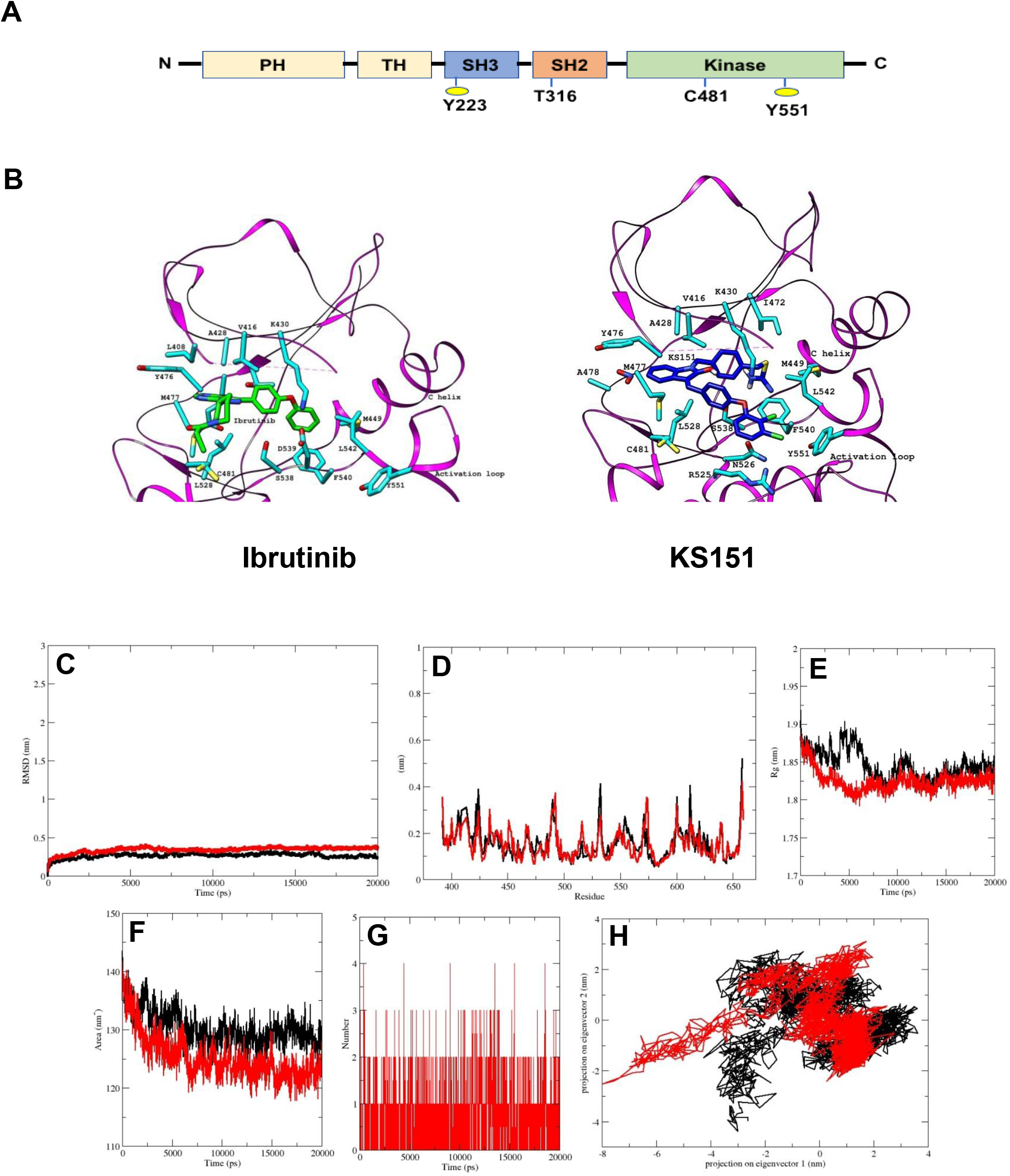
KS151 interacts with the BTK. ***A***, structural domain of BTK. BTK contains five domains, which includes the amino terminal pleckstrin homology (PH) domain, a proline-rich TH domain, SRC homology (SH) domains SH2 and SH3, as well as the kinase domain. ***B***, the interaction of ibrutinib and KS151 to the activation loop of BTK. On the left, the crystal structure of BTK bound to Ibrutinib and on the right the crystal structure of BTK protein bound with KS151. ***C-H***, conformational stability of hBTK (black) and hBTK-KS151 complex (red) during 20ns molecular docking simulation **(C),** Backbone RMSD differences of hBTK-apo and hBTK-KS151 complex (**D**) RMSF differences hBTK-apo and hBTK-KS151 complex (**E**) Radius of gyration differences between hBTK-Apo and hBTK-KS151 complex (**F**) Average number of H-bonds as a function of time in hBTK-Apo and hBTK-KS151 complex (red). **(G)** Accessible surface area in hBTK-Apo and hBTK-KS151 complex. **(H)** Projection of C_α_ atoms in essential subspace along the first two eigenvectors of hBTK (black) and hBTK-KS151 complex (red) showed different projection spaces.

Development of KS151 started with our first-generation BTK inhibitor, KS99 [47]. KS99 suppressed *in vitro* kinase activities of BTK and its phosphorylation at Tyr 223 in MM U266 cells [47]. We further demonstrated the cytotoxic potential of KS99 in CD138^+^ and CD138^neg^ (stem cell-like) cells isolated from myeloma patients [47]. Interestingly, bortezomib failed to show any response in these cells [47]. Because ibrutinib was found to be ineffective due to the mutation at C481, we sought to modify KS99 in such a way that it could avoid this residue. Our study demonstrates that KS151 is a specific inhibitor of BTK and is more potent than ibrutinib and other chemotherapeutic agents, effective against bortezomib resistant cells, potentiates the response of bortezomib, lenalidomide, and panobinostate in stem cell-like cells, and regulates the expression of stem genes.

## MATERIALS AND METHODS

### Reagents

RPMI 1640, fetal bovine serum (FBS), 0.4% trypan blue vital stain, and an antibiotic–antimycotic mixture were obtained from Life Technologies (Carlsbad, CA). Rabbit anti-phospho-BTK (#5082), rabbit anti-BTK (#8547), rabbit anti-phospho-PLCγ2 (#3871), rabbit anti-PLCγ2 (#3872) were obtained from Cell Signaling Technology (Danvers, MA). Antibodies against GAPDH, Hoechst 33342, 3-(4,5-dimethylthiazol-2-yl)-2,5-diphenyltetrazolium bromide (MTT), Goat anti-rabbit and goat anti-mouse horseradish peroxidase conjugates were purchased from GE Healthcare (Piscataway, NJ) and goat anti-rabbit Alexa Fluor 594 was purchased from Molecular Probes (Eugene, OR). The ADP-Glo Kinase assay kit along with BTK enzyme was procured from Promega Corporation (Madison, WI, USA). Cell viability dye Viva Fix 498/521 (#1351115) and mouse anti-human CD138 Alexa Fluor 647 (#MCA2459A647) were obtained from Bio-Rad (Hercules, California). CD138 magnetic beads were procured from Miltenyi Biotec (Auburn, California). The TaqMan primers specific for GAPDH (Hs02786624_g1), BTK (Hs00975865_m1), Nanog (Hs02387400_g1), Gli1(Hs00171790_m1), and Myc (Hs00153408_m1) were obtained from Life Technologies (Carlsbad, CA). KS151 was synthesized and characterized at Penn State College of Medicine, Hershey, PA. A 25 mmol/L solution of KS151 was prepared in dimethyl sulfoxide, stored as small aliquots at −20°C, and then diluted as needed in cell culture medium.

### Cell culture

Human MM cells MM.1R, MM. 1S, U266, RPMI 8226 cells, and BM stromal cells (BMSCs) were purchased from the American Type Culture Collection (ATCC) (Manassas, VA). Bortezomib resistant cells MM.1S BTZ-R and RPMI8226 BTZ-R were a generous gift from Dr. Nathan Dolloff, The Medical University of South Carolina, Charleston, SC, USA. MM.1R, MM. 1S, and RPMI 8226 cells were cultured in RPMI 1640 growth media (Cellgro, Manassas, VA) supplemented with heat-inactivated 10% Fetal Bovine Serum (Sigma, St. Louis, MO), 100 I.U./ml penicillin, and 100 μg/ml streptomycin (Cellgro, Manassas, VA). U266 cells were cultured in RPMI 1640 with 15% FBS. BMSCs were procured from ATCC and cultured in Dulbecco’s Modified Eagle Medium (DMEM) (ATCC) supplemented with heat-inactivated 10% Fetal Bovine Serum (Sigma), 100 I.U./ml penicillin, and 100 μg/ml streptomycin (Cellgro, Manassas, VA). All cells were cultured in a 37°C and 5% CO2 incubator.

### Isolation of CD138^neg^ cells

CD138^neg^ cells were isolated by two methods using the S3e ^TM^ Bio-Rad Cell Sorter and magnetic beads. MM.1R, and RPMI 8226 cells were washed and resuspended in cold PBS with 3% FBS. Then cells were stained with cell viability dye VivaFix498/521 and mouse anti-human CD138 Alexa Fluor 647 and incubated in the dark on ice for 25 minutes. After washing twice, cells were sorted using S3e ^TM^ Bio-Rad following the manufacturer’s protocol [48]. Alternatively, CD138 magnetic beads were used to isolate CD138^neg^ cells. Human MM cells MM.1R and RPMI 8226 cells were fractioned by a MACS separator, using CD138 magnetic beads. After washing cells from the media, cells were resuspended in 0.5% MACS buffer and incubated with CD138 microbeads at 4°C for 15 minutes. After washing, CD138^neg^ cells were isolated using an MS column according to the manufacturer’s protocol [47].

### Protein and ligand preparation

The three-dimensional crystal structure of the human Bruton’s tyrosine kinase hBTK (PDB: 3GEN) receptor was retrieved from the Protein Data Bank and used as a receptor for docking [49]. In order to prepare the protein for docking, protein structure was prepared by stripping off water molecules and other bound heteroatoms from structure coordinates file using PyMOL software and hydrogen atoms were added into the receptor by using MGL Tools and converted in PDBQT format for further analysis [50, 51]. Chem Draw 20.0 was used to create the 3D structures of analogs in mol2 format, which was then converted into protein data bank (PDB) format using PyMOL software.

### Molecular docking

Molecular docking aids in predicting a ligand’s preferred binding pose with a protein and analyzing inhibitory interactions between them. Here a, molecular docking studies were performed using Autodock Vina [51]. A variety of searches were conducted in order to provide the best possible conformation. The docking grid size in X, Y, Z dimensions was set at size_x = 50.00 A^0^, size_y = 50.00 A^0^ and size_z = 50.00 A^0^, respectively and centered at center_ x = - 17.03, center_y = 10.28 and center_ z = 21.22 with a 0.375 A^0^ grid spacing and exhaustiveness were set to 24. KS151 was prepared by setting the active torsions and detecting the root of the molecules to adjust according to the defined pocket of the target protein. The docking studies were carried out with rigid protein and the molecular docking outcomes were analyzed based on binding energy (kcal/mol) and binding interactions profile. The schematic diagrams of protein ligand interactions were generated using UCSF Chimera (https://doi.org/10.1002/jcc.20084) and analyzed using Ligplot+ v.2.1 [52].

### Molecular dynamics simulation

The MD simulations were carried out up to 20 ns to analyze the stability of the hBTK-KS151 complex using GROMACS 2018.1 running on an Intel(R) Core (TM) i5-9400 CPU machine installed with Ubuntu 18.04 Linux. In all runs, the GROMOS43a1 force field was used [53]. The PRODRG server was used to generate ligand topology files [54]. Before simulation, both the hBTK and hBTK-KS151 complexes were solvated in the TIP3P model in a triclinic box with a minimum spacing of 2.5 Å distance between any protein atoms to the closest box edge. To neutralize the system Na^+^ ions were added, and 50000 steps of steepest descent energy minimization were performed. Systems were equilibrated using NVT and NPT ensembles for 100 ps at 300K and 1 bar of pressure using a modified Berendsen thermostat (tau_T = 0.1 ps) and a Parrinello–Rahman barostat (tau_P = 2 ps). The final simulation run lasted for 20 ns, and the dynamic trajectories of the root mean square deviation (RMSD), the root mean square fluctuation (RMSF), radius of gyration (Rg), solvent accessible surface area (SASA), hydrogen bonding, and projections of C_α_ atoms were visualized by the XMGRACE software package [55].

### Synthesis of KS151

The chemicals and solvents were purchased from commercial vendors. Reactions were carried out using dried glassware under an atmosphere of nitrogen. Reaction progress was monitored with analytical thin-layer chromatography (TLC) on aluminum backed precoated silica gel 60 F254 plates (E. Merck). Column chromatography was carried out using silica gel 60 (230-400 mesh, E. Merck) with the solvent system indicated in the procedures. Nuclear magnetic resonance (NMR) spectra were recorded using a Bruker Avance II 500 spectrometer.

KS151 was synthesized as shown in **Figure 2**. Briefly, to a solution of 4-iodobenzaldehyde (**1**) (7.5 mmol) in dioxane (30 ml) was added 2,3-dichlorophenol (**2**) (7.5 mmol), cuprous iodide (0.75 mmol), N, N-dimethylglycine hydrochloride (2.25 mmol), and cesium carbonate (15 mmol). The mixture was stirred at 105°C in a nitrogen atmosphere for 18h. The mixture was concentrated and extracted with ethyl acetate, washed with water, and dried over MgSO4. The crude product was purified by a silica gel column using hexane and ethyl acetate (9:1) as the eluent to yield the intermediate (**3**). Compound (**3)** (3.37 mmol) was refluxed with 5-Nitro-2-oxindole (3.37 mmol) and piperidine in ethanol for 8 h and filtered to obtain the product (**4**). Compound (**4**) (1.75 mmol) was dissolved in anhydrous DMF (30 mL), and K2CO3 (2.2 mmol) was added and stirred for 1h. 1,4-Bis(bromomethyl)benzene (7 mmol) was added slowly with constant stirring until compound **4** had been consumed (monitored by TLC). The reaction mixture was poured into cold water and extracted with ethyl acetate. The ethyl acetate layer was washed with water and brine and dried over MgSO4. The solvent was removed by rotavaporation, and the crude product was purified by silica gel column chromatography using hexanes/ ethyl acetate (80:20) as the eluent to yield the intermediate (**5**) as yellow crystals. To compound **5** (1.04 mmol), thiourea (1.0 mmol) and ethanol (25 mL) were added and heated to reflux until the starting material had been consumed (monitored by TLC). The solvent was removed under a vacuum. The final product 4-((3-(4-(2,3-dichlorophenoxy) benzylidene)-5-nitro-2-oxoindolin-1-yl) methyl) benzyl carbamimidothioate hydrobromide (**6**) (**KS151**) was recrystallized in ethanol-ethyl acetate to afford **KS151** (yield 70%). The identity of **KS151** was confirmed by nuclear magnetic resonance and mass spectral analysis, and purity (>99%) was quantified by high-performance liquid chromatography analysis. Yellow solid, yield: 70%; ^1^H NMR (500 MHz, DMSO-d6): δ 9.17 (s, 2H), 8.98 (s, 2H), 8.8 (d, J = 2.5 Hz, 1H), 8.6 (d, J =9.0 Hz, 1H), 8.41 (d, J = 2.0 Hz, 1H), 8.32 (m, 4H), 8.01 (s, 1H), 7.89 (d, J = 8 Hz, 1H), 7.12-7.60 (m, 6H), 5.10 (s, 2H), 4.45 (s, 2H); MS (ESI) m/z 687 [M+H].

**Figure 2:**
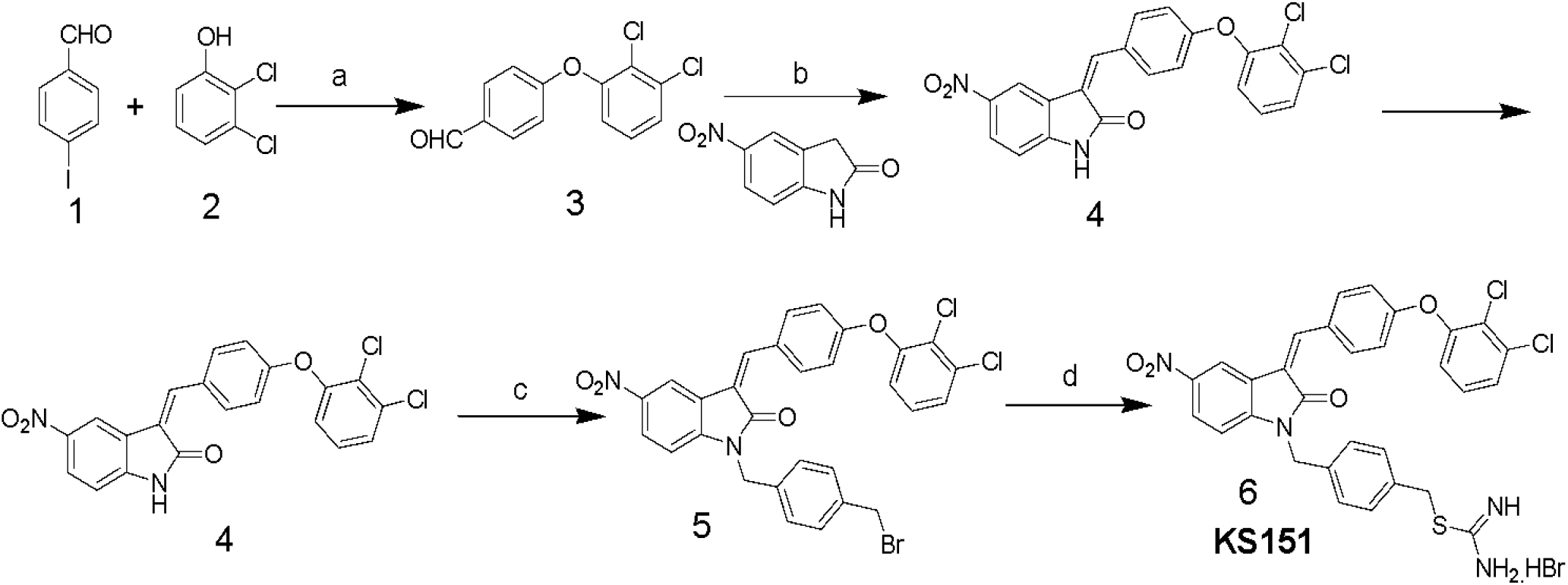
Scheme of KS151 synthesis and conditions: a) Cs_2_CO_3_, Cul, N,N-dimethylglycine, Dioxane; b) EtOH, pipiridine, reflux, 8h; c) K_2_CO_3_/DMF, BrCH_2_C_6_H_4_CH_2_Br, 8h; and d) Thiourea, EtOH, 100°C, 4h.

### *In vitro* kinase assay

The assay was performed as described earlier [47]. Various concentrations of KS151 (0.5 – 1μM) were used in the reaction. Briefly, various amounts of KS151 were diluted in kinase assay buffer consisting of BTK enzyme, substrate, and adenosine triphosphate (ATP) (total volume, 5 mL). The assay was performed in a 96-well plate. The reaction mixture was incubated at room temperature for 1h. After incubation, 5 μL of ADP-Glo reagent was added, and the reaction was incubated for an additional 40 min at room temperature. Adding 10 μL of kinase detecting reagent, followed by 30 min of room temperature incubation, terminated the reaction. Finally, luminescence was detected using an Infinite M200 Pro multi plate reader (Tecan, Switzerland). Kinase activity was normalized by vehicle control (dimethylsulfoxide, DMSO) and a graph was performed using Prism version 5 software (GraphPad, La Jolla, CA).

### RNA extraction, reverse transcription, and qPCR analysis

RNA was extracted from cells using an RNA extraction kit (Invitrogen) and converted to cDNA using an Invitrogen kit (#12183018A) as per the manufacturer’s instructions [47]. RNA was quantified using an ND-1000 spectrophotometer (Nanodrop Technologies, Wilmington, DE). Reverse transcription (RT) was done to obtain complementary DNA, and samples were diluted to prepare for qPCR as previously described [47, 56]. Quantitative polymerase chain reaction (qPCR) experiments were performed using an AB Applied Biosystems Step One Pulse Real-Time PCR System and TaqMan Universal PCR Master Mix (Applied Biosystems, Foster City, CA). The housekeeping gene GAPDH was used for all delta-delta Ct calculations as described earlier [57].

### Western blot

Human MM cells were treated with KS151 for 24h. After incubation, cells were washed and lysed using RIPA buffer supplemented with protease and phosphatase inhibitor cocktails [58]. After incubation on ice for 1h, cell extracts were vortexed and the cell lysate was centrifuged at 15,000 rpm for 10 minutes. The supernatant was kept at –80^0^C until further use. Protein was quantified using a BCA Protein Assay kit (Pierce Scientific, Rockford, IL). Heat denatured protein samples were separated using a 4–12% NuPAGE gel (Thermo Scientific, Rockford, IL, USA). Proteins were transferred to PVDF membranes at 30 volts for 90 minutes. After blocking with 5% non-fat dry milk for 1h, membranes were incubated overnight at 4^0^C with primary antibodies. After washing, the secondary antibody was added and incubated for 3h at room temperature. After washing, the membrane was exposed to an enhanced chemiluminescent substrate for detection of horseradish peroxidase (HRP) (PierceTM ECL Western; Thermo Scientific. Rockford, IL, USA). Bands were visualized and quantitated using the Chemi DocTM MP imaging system (Bio-Rad, Hercules, CA).

### MTT assay

The effect of KS151 and the other drugs on cell viability was determined using an MTT assay by a method described earlier [47]. Briefly, 5000 cells of human MM were plated in triplicate in 96-well plates and treated with various treatments for 72h [47]. After incubation for 72h, 20 μl of a freshly prepared MTT solution (5 mg/mL) was added and incubated for 3h at 37^0^C. The 96-well plates were centrifuged, the supernatant was removed, and 100 μl of dimethyl sulfoxide (DMSO) was added to each well to dissolve the formazan crystals. The optical density of the resulting solution was measured by delta value (570-630 nm) using a 96-well multi-mode microplate reader (BioTek Technologies, Winooski, VT, USA) [47].

### Annexin V live dead assay

To study apoptosis, a Muse® Cell Analyzer (Luminex Corporation, Austin) was used. MM cells were treated with either KS151 or in combination with other drugs for 24h before subjected to a Luminex Annexin V Live Dead Assay (Luminex Corporation, Austin). Annexin V live dead assay was performed essentially following the manufacturer’s protocol. Briefly, cells were plated at a density of 2×10^5^ per well and then treated and incubated for 24h. Then, cells were scraped and mixed with Annexin V stain (Luminex Corporation) and incubated for 20 minutes. The Muse Cell Analyzer was used to analyze the data. The lower left quadrant [annexin V neg and 7-amino-actinomycin neg (7-AAD)] reflects the live cells, while the lower right quadrant (annexin V+ and 7-AAD neg) contains early apoptotic cells. The late apoptotic and dead cells are in the upper right quadrant (annexin V+ and 7-AAD+). The upper left quadrant represents the dead cells (annexin V and 7-AAD+).

### Immunostaining and confocal microscopy

Cells were plated in 12 well plates and treated with 5 μM of KS151 for 24h. Then 100 μl of cell suspension was loaded into the funnel that is used for the Cytospin. Cells were centrifuged for 3 minutes at 800 rpm, followed by fixing with and 4% formaldehyde for 15 minutes. After washing three times with TBST, the cells were incubated with 0.2 % Triton while rocking for 18 minutes. Then, after 1h of blocking with 3% BSA, 1:200 primary antibody was added in 3% BSA overnight at 4^0^C. After washing three times with phosphate buffered saline (PBS), secondary antibody, goat anti-rabbit IgG (H+L) Alexa Fluor ™ Plus 488 (Red A32731, Invitrogen) was added at a 1:200 dilution and kept rocking for 1h at room temperature. Slides were mounted with ProLong ^TM^ Diamond Antifade Mount with DAPI (# P36971, Invitrogen, Carlsbad, CA) and covered with a coverslip, then imaged with the Nikon A1R GaAsP Laser Scanning Confocal Microscope (Nikon Eclipse Ti inverted using a 60x objective). The Integrated density of cells was calculated using the ImageJ software as previously described [59].

### Co-culture of MM and BM stromal cells

A Co-culture of MM cells with bone marrow stromal cells (BMSCs) was performed by a method described before [60]. In brief, 2×10^5^ BMSCs were seeded in six well plates overnight. After incubation, 2×10^5^ MM cell lines were plated on top of BMSCs and treated with 5 μM of KS151 for 24h. The cells were collected using a scraper, and RNA was harvested, and qPCR was performed as described above.

### Statistical analysis

Statistical significance of the results was analyzed by the Student’s t-two tail test using GraphPad Prism software.

## RESULTS

### *In silico* molecular docking and molecular dynamics simulations of KS151

First, we performed *in silico* analysis using the three-dimensional crystal structure of the human BTK (PDB: 3GEN) retrieved from the Protein Data Bank [49]. The phenoxyphenyl group of isatin of KS99 was overlaid with the same fragment of ibrutinib, which completely entered the hydrophobic pocket and formed a face-to-edge π-stacking interaction with surrounding amino acid residues. The central core of ibrutinib with 4-aminopyrazolo [3,4-d] pyrimidine maintains the potency of inhibiting BTK; we selected isatin as the new scaffold to replace the pharmacophore of ibrutinib by employing the theory of bioisostere. *In silico* analysis, illustrated in **Figure 1B**, revealed KS151 docked with hBTK, showed potential binding against hBTK with a docking score of −10.3 kcal/mol. KS151 forms backbone peptide bonds of residues Met477 and Ala478 at a distance of 3.20 A^0^ and 3.33 A^0^, respectively. Amino acid residues, Leu408, Val416, Met449, Leu528, and Phe540 were involved in hydrophobic interactions while Thr474, Tyr476, Asn526 and Asp539 are in close proximity to create hydrophilic-environment within the active site pocket. The 1,4-bis (bromomethyl) benzene was selected as a linker to connect the scaffold and isothiourea warheads.

To further probe the stability and dynamics of the hBTK-KS151 complex in active site pocket, a molecular dynamics simulation calculation was carried out for 20 ns. The trajectories of hBTK and hBTK-KS151 complex were studied to understand the flexibility and stability of the ligand complex. Root mean square displacements (RRMSD), root mean square fluctuations (RMSF), radius of gyration (Rg), and hydrogen bond contact were evaluated as a function of time. The average RMSD values of hBTK alone and the KS151 complex are 0.23 nm 0.37 nm respectively. During the MD simulation, the overall residual motions of the docked complex were computed. The RMS fluctuations for the amino acid residues of the hBTK and KS151 complex are in the same pattern without any major deviations. Due to ligand binding, it is not highly reflective. The RMSF plot indicated stable interactions in the binding pocket throughout the simulations. The radius of gyration (Rg) of a protein is a measure of its compactness. If a protein is stably folded, its Rg value is likely to remain stable. If a protein unfolds, its Rg will change over time. Radius of gyration values for the complex were 1.84 nm compared to 1.86 nm of hBTK indicating a gain in compactness upon ligand binding. The average SASA value of the complex was estimated at 121.9 compared to 129.7 for hBTK alone. The intermolecular H-bonding between KS151 and hBTK is critical for the complex’s stability. Hydrogen bonding interactions between KS151 and various residues in binding pocket ranges from 3 to 4 at different time intervals of the KS151 simulated MD trajectories indicating stable interaction patterns. The results of these analyses are summarized in **Figure 1C-H.**

### KS151, a specific inhibitor of BTK, inhibits in vitro kinase activities

We synthesized sixteen analogs by following the scheme shown in **Figure 2**. All analogs were purified by silica gel column and purity was determined by analytical high-pressure liquid chromatography followed by NMR and mass spectrometry. The efficacies of new analogs were tested against BTK by performing an *in vitro* kinase assay, which allowed us to identify KS151 as a lead molecule. Indeed, KS151 inhibited 44% and 49.8% of kinase activity at 0.5 and 1 μM respectively, whereas ibrutinib inhibited 25.9% and 32.2% of kinase activity at 0.5 and 1 μM respectively **(Figure 3A)**. This suggests that KS151 is more potent than ibrutinib. We next determined whether KS151 is a selective BTK inhibitor. We tested the specificity of KS151 by performing a semi-quantitative, sandwich-based array following the manufacturer’s protocol (RayBiotech Life, Peachtree Corners, GA, USA) for detecting 71 tyrosine kinase phosphorylation signatures. We detected only 23 kinases in the human MM cell line U266 **(Figure 3B)**. Among 23 detectable kinases, we observed that KS151 specifically inhibits BTK. This suggests that KS151 selectively targets BTK cells and has little or no effect on the cell kinases that were tested.

**Figure 3:**
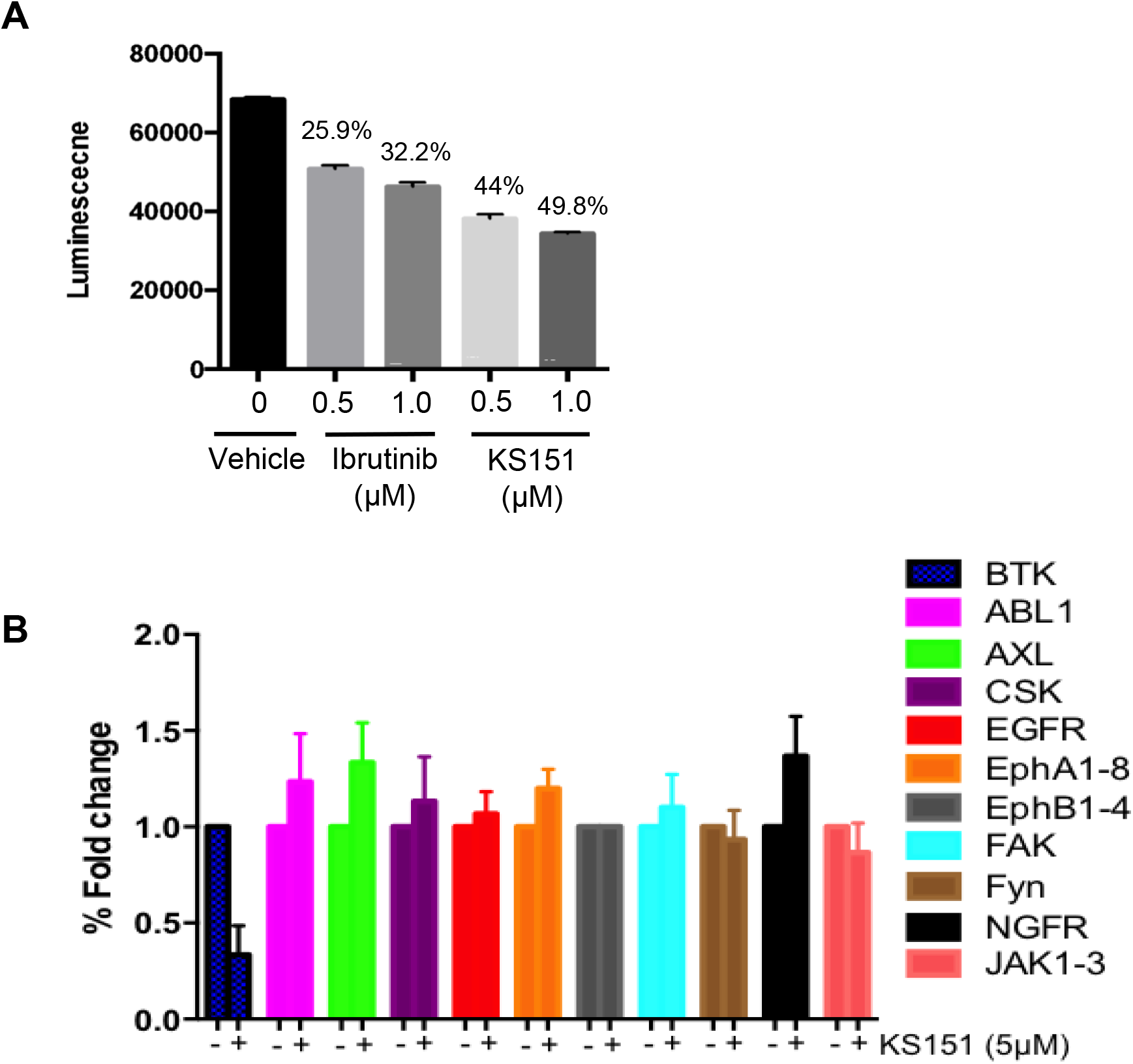
KS151 inhibits the *in vitro* BTK kinase activities and specific to BTK. ***A***, KS151 suppresses *in vitro* kinase activities in a dose-dependent manner. Different doses of KS151 were used to determine the kinase inhibitory response. Values written on top of bars indicate the percentage inhibition of kinase activities. ***B***, the specificity of KS151 was assessed by a semi-quantitative, sandwich-based array for 71-tyrosine kinase phosphorylation signatures. The RayBio Human RTK Phosphorylation Antibody Array kit was used to investigate the effect of KS151 on various kinases. Data represents only those expressed kinases in MM cells.

### KS151 inhibits BTK signaling pathways

To choose the best cell line model to study the effect of KS151, we tested various MM cell lines for BTK expression. We extracted proteins from untreated U266, MM. 1R, MM. 1S, and RPMI 8226 MM cell lines and investigated BTK expression using western blot **(Figure 4A).** We found that RPMI 8226 has the highest level of endogenous BTK followed by MM.1S, MM.1R, and with U266 having the least expression. The expression of the mRNA of BTK was also assessed in each cell line. Similar to the western blot results, we found that RPMI 8226 has the highest BTK expression followed by MM.1S, and MM. 1R **(Figure 4B).** We further investigated whether KS151 inhibits the phosphorylation of BTK and its downstream target PLCγ2. The autophosphorylation of BTK at Y223 was inhibited by KS151 in a dose-dependent manner. Furthermore, KS151 inhibited phospho-PLCγ2 without affecting the PLCγ2 **(Figure 4C)**. We also used confocal microscopy to test the effect of KS151 on phospho-BTK. We stained fixed MM cells for phospho-BTK after treating cells with 5 μM KS151 for 24h. The total signal intensity was calculated. We found that KS151-treated cells had a lower integrated density of phospho-BTK compared to the untreated control **(Figure 4 D-E).** We calculated the median reading of integrated density. Interestingly, 5 μM of KS151 was able to inhibit phospho-BTK by 56.48% in the RPMI 8226 cell line. Interestingly, treatment of KS151 inhibits the nuclear translocation of BTK. It is important to mention that BTK translocation in different compartments has been observed [61, 62].

**Figure 4:**
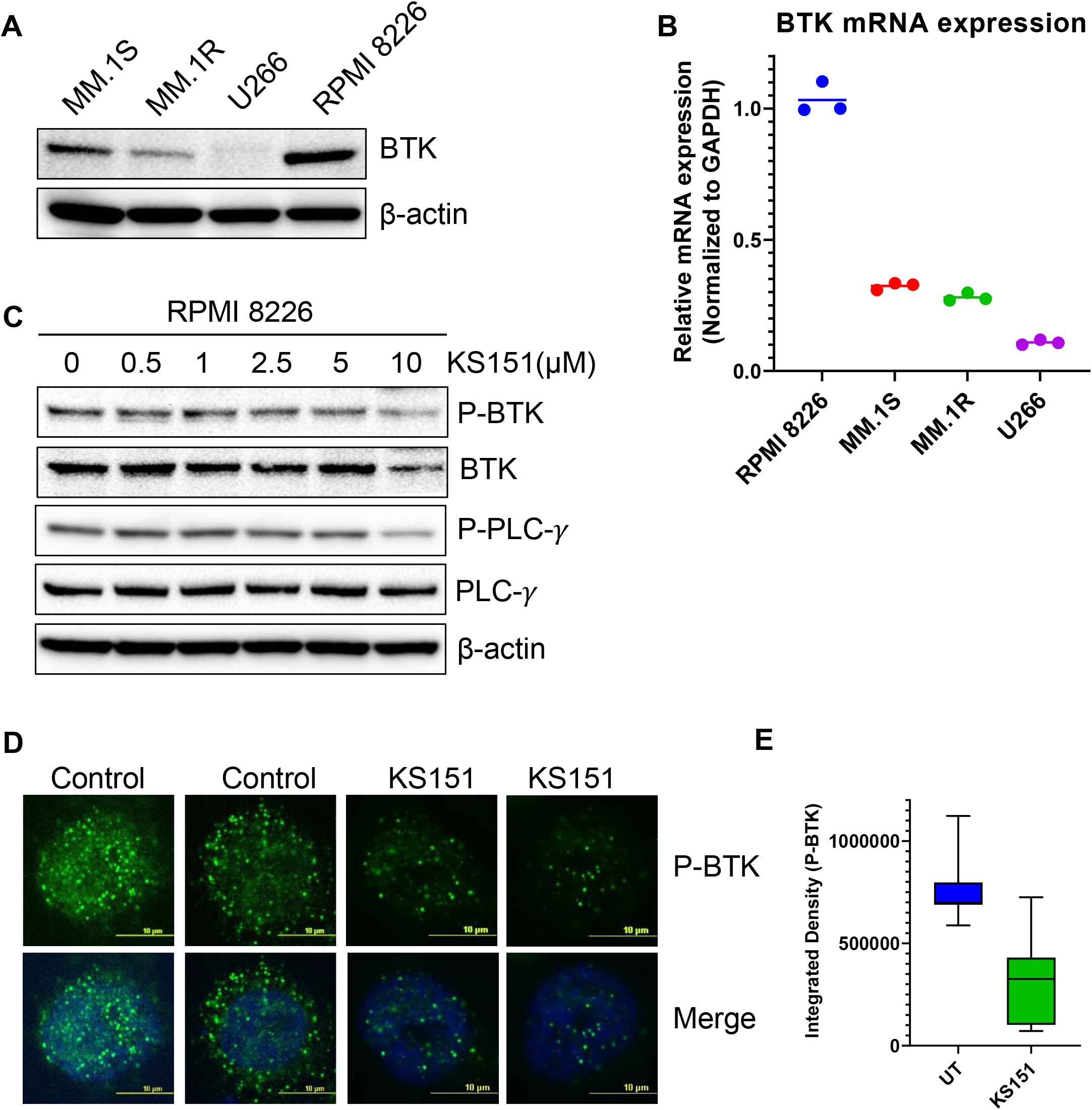
KS151 inhibits BTK signaling pathways. ***A***, Expression of BTK in different MM cell lines determined by western blot. ***B***, Expression of BTK in various MM cell lines determined by real-time quantitative PCR (QPCR). GAPDH was used as an internal control. ***C***, KS151 suppresses phosphorylation of BTK and PLCγ2 in a dose-dependent manner. MM.1R cells were treated with mentioned amount of KS151 for 24h and immunoblotting was performed. As a loading control, the stripped membrane was probed with either BTK, or PLCγ2 or β-actin antibodies. ***D***, A confocal microscopy representative for the effect of 5 μM KS151 on phospho-BTK after 24h treatment. ***E***, Mean signal intensity calculated via Nikon software to compare phosphorylation between KS151 treated and untreated control.

### KS151 is cytotoxic against naïve and Bortezomib resistant MM cells and more potent than other chemotherapeutic agents

Once the specificity and functionality of KS151 was established, we tested its cytotoxic potential and compared it with existing chemotherapeutic agents via the MTT assay [47]. MM cell lines MM.1S, MM.1R, U266, and RPMI 8226 were treated with various concentrations of KS151 (0.1-25 μM) for 72h. An MTT cell viability assay confirmed a dose-dependent decrease in cell viability of MM cell lines by KS151, which inhibited the growth of MM.1S, MM.1R, U266, and RPMI 8226 with an IC50 value of 12.07, 8.4, 15.48, and 9.07 μM, respectively **(Figure 5A)**. Furthermore, we compared its effect to chemotherapeutic compounds that are FDA-approved to treat MM. We compared KS151 to different classes of chemotherapeutic agents such as BTK inhibitors (acalabrutinib, and ibrutinib), alkylating agents (cyclophosphamide, and melphalan), immunomodulatory agents (lenalidomide), histone deacetylase (HDAC) inhibitor (panobinostat), and plant alkaloid “topoisomerase II inhibitor” (etoposide). Interestingly, we found KS151 more potent than most of these medications in MM.1R and RPMI 8226 MM cell lines **(Figure 5B)**. Because the acquired resistance against bortezomib is the main reason for relapse in MM, and we further tested the extent to which KS151 is effective in bortezomib-resistant cells. We compared the efficacy of KS151 with other BTK inhibitors (acalabrutinib, and ibrutinib) in bortezomib resistant cell lines. MM.1S-bortezomib-resistant (MM.1S Btz-R), and RPMI 8226-bortezomib-resistant (RPMI 8226-Btz-R) cell lines were treated with various concentrations of KS151 (0.1-25 μM) for 72h. As reflected in **Figure 5C**, KS151 was found to be more effective in these resistant cells than acalabrutinib, and ibrutinib. The IC50 value of KS151 was 7.38 μM, and 5.13 μM in MM.1S Btz-R, and RPMI 8226-Btz-R cell lines respectively, whereas in comparison, IC50 was much higher for ibrutinib (16.7 μM, 15.28 μM) and acalabrutinib (more than 25 μM) in both cell lines **(Figure 5C)**. To investigate whether the decrease in MM cell viability after KS151 treatment was due to apoptosis induction, we assessed the apoptosis-inducing potential of KS151 in MM cells by Annexin V + 7AAD staining. Various MM cells were treated with increasing concentrations of KS151 for 24h, and apoptotic assay was performed using the Muse Cell Analyzer. KS151 induced the apoptosis in a dose-dependent manner in various MM cell lines (**Figure 5 D-E**).

**Figure 5:**
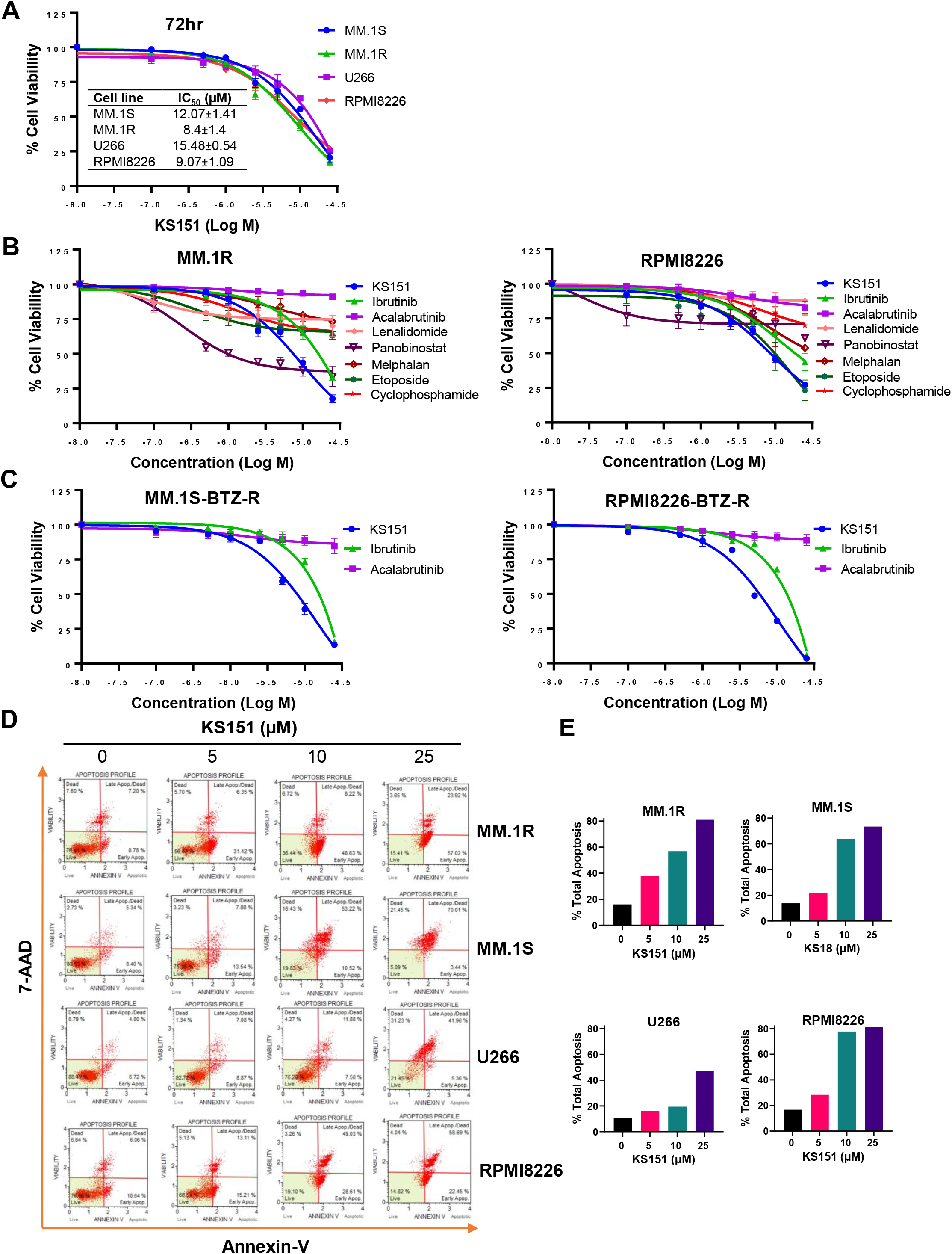
The cytotoxic effect of KS151 on MM and bortezomib -resistant cells. ***A***, Various MM cells were treated with increasing amount (0.1-25 μM) of KS151 for 72h and cell viability was assessed by MTT as described in the materials and methods section. IC50 was calculated by a non-linear regression plot. ***B***, MTT cell viability assay investigating the proliferation of MM.1R and RPMI 8226 MM cell lines treated with different doses (0.1-25 μM) of different chemotherapeutic agents (acalabrutinib, ibrutinib, lenalidomide, panobinostat, melphalan, etoposide, cyclophosphamide) as well as KS151 for 72h. The cell viability was assessed by MTT as described in the materials and methods section. ***C***, Bortezomib-resistant MM cells were treated with increasing amount (0.1-25 μM) of KS151, ibrutinib, and acalabrutinib for 72h and cell viability was assessed by MTT. ***D***, Annexin V staining to detect apoptotic cells. MM.1S, MM.1R, U266, and RPMI 8226 MM cells treated for 24h with different doses of KS151 and then stained with Annexin V dye and analyzed by the Muse Cell Analyzer. ***E***, Quantitation of apoptotic cells (early plus late apoptosis) after treatment as detected by Annexin V staining.

### KS151 potentiates the cytotoxic effect of Bortezomib, Lenalidomide and Panobinostat in both MM and MM stem cell-like cells

Because bortezomib, lenalidomide, and panobinostat are widely used to treat MM, we first investigated the extent to which KS151 augments the efficacy of therapeutic drugs in MM cell lines. We assessed the apoptosis-inducing potential of KS151 in combination with bortezomib, lenalidomide, and panobinostat on MM cells by Annexin V + 7AAD staining. As shown in **Figure 6A,** the addition of KS151 to chemotherapeutic drugs had minimal effect as bortezomib, lenalidomide and panobinostat induced apoptosis even alone in MM.1R cells. However, in RPMI 8226 cells that contain a higher expression of BTK, the addition of KS151 to therapeutic agents potentiates the apoptotic response **(Figure 6B),** suggesting that BTK inhibition is required to improve the therapeutic outcome of bortezomib, lenalidomide, and panobinostat. As reflected from **Figure 6A&B**, although current treatments may kill the tumor bulk, they can leave behind therapy-resistant stem cell-like population, which serves as a reservoir for disease recurrence [63, 64]. Therefore, approaches to target stem cell-like cells to prevent cancer resistance are important for managing MM [6, 64]. Our goal was to test whether KS151 compromises the survival of stem cell-like cells. We isolated CD138^neg^ cells (which possess stem cell-like properties as discussed above) from human MM.1R, and the RPMI 8226 cell line and treated them with KS151 using a cell sorter. We found that KS151 induced the apoptosis in both CD138^neg^ MM.1R, and CD138^neg^ RPMI 8226 cells **(Figure 6C).** We further tested whether a combination of KS151 with bortezomib, lenalidomide, or panobinostat is an effective strategy against CD138^neg^ cells. We treated CD138^neg^ MM.1R, and CD138^neg^ RPMI 8226 cells with various concentrations of KS151. The combination of KS151 further potentiates the response of bortezomib, lenalidomide, and panobinostat, suggesting that KS151-based treatment combinations are effective against stem cell-like cells, which are the main source of MM relapse **(Figure 6D-E)**.

**Figure 6:**
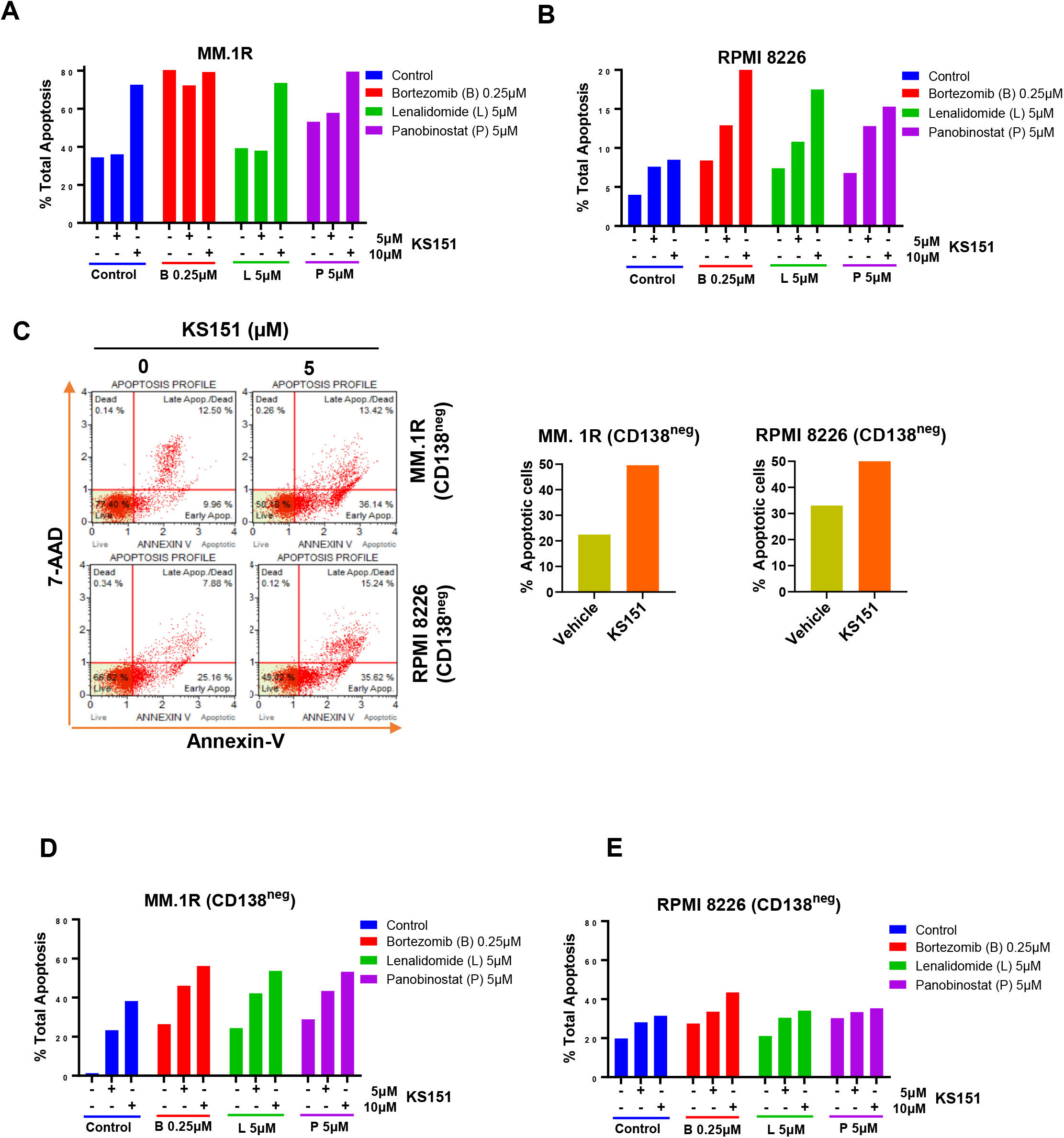
Combination studies of KS151 with other chemotherapeutic agents. ***A-B***, Annexin V staining to detect either MM.1R or RPMI 8226 apoptotic cells. MM.1R or RPMI 8226 cells were treated with 5 or 10 μM of KS151 alone or in combination with 0.25 μM bortezomib, 5 μM lenalidomide, or 5 μM panobinostat. After 24h, cells were stained with Annexin V dye and analyzed by the Muse Cell Analyzer by a method described in the materials and methods section. **C**, CD138^neg^ cells of MM.1R and RPMI 8226 were isolated using magnetic beads and treated for 24h with KS151. Cells were stained with Annexin V dye and analyzed by Muse Cell Analyzer. ***D-E***, Annexin V staining to detect CD138^neg^ MM.1R and CD138^neg^ RPMI 8226 apoptotic cells. After sorting CD138^neg^ cells from MM.1R or RPMI 8226 with Bio-Rad cell sorter, cells were kept in the incubator overnight to recover and then treated with 5 or 10 μM of KS151 alone or in combination with 0.25 μM bortezomib, 5 μM lenalidomide, or 5 μM panobinostat. After 24h, cells were stained with Annexin V dye and analyzed by Muse Cell Analyzer.

### KS151 inhibits the stemness markers

BTK regulates the expression of stem genes (OCT4, SOX2, Nanog, and MYC) [17, 18]. Given our results in **Figure 6C-E**, we examined if KS151 impacts the expression of stem genes in stem cell-like cells (CD138^neg^). Therefore, we investigated whether KS151 can also inhibit stemness markers in CD138^neg^ MM.1R and CD138^neg^ RPMI 8226 cells. MM.1R CD138^neg^ and CD138^neg^ RPMI 8226 cells were isolated from MM.1R using magnetic beads. Then, cells were treated with 5 μM of KS151. After 24h, RNA was isolated from cells, and RT reaction and qPCR were performed. We found that KS151 inhibited the stemness markers effectively in both stem cell-like cells. The stemness marker Gli1, Myc, and SOX2 were inhibited by 25.6, 31.6., and 74.9% respectively in CD138^neg^ MM.1R cells **(Figure 7B)**. Similarly, the stemness marker Gli1, Myc, and SOX2 were inhibited by 44, 32, and 13.9% respectively in CD138^neg^ RPMI 8226 cells. Altogether, the results in **Figure 7 A-B** suggest that KS151 targets MM cells with stem cell-like properties. It has been demonstrated that BM microenvironment (BMM) regulates the stemness of MM cells via BTK signaling pathways [18]. To examine the efficacy of KS151 in the relevant microenvironmental context, we tested the efficacy of KS151 when MM cells are co-cultured with BMSCs. We co-cultured MM.1R and RPMI 8226 cells on top of BMSCs in a 1:1 ratio. Then, cells were treated with 5 μM of KS151. After 24 hours, RNA was isolated from cells and RT-qPCR was performed using specific primers for Nanog, and Gli1. As shown in **Figures 7 C-D,** KS151 was able to suppress stemness markers even in the presence of BMSCs. Nanog and Gli1 were inhibited 67.48%, and 46.2% in MM.1R co-culture, respectively. Likewise, Nanog and Gli1 were inhibited by 24.8%, and 56.5%, respectively, in RPMI 8226 and BMSCs co-culture, suggesting that novel BTK inhibitor regulates the stem cell-like phenotypes by modulating the microenvironment **(Figure 7 C-D)**.

**Figure 7:**
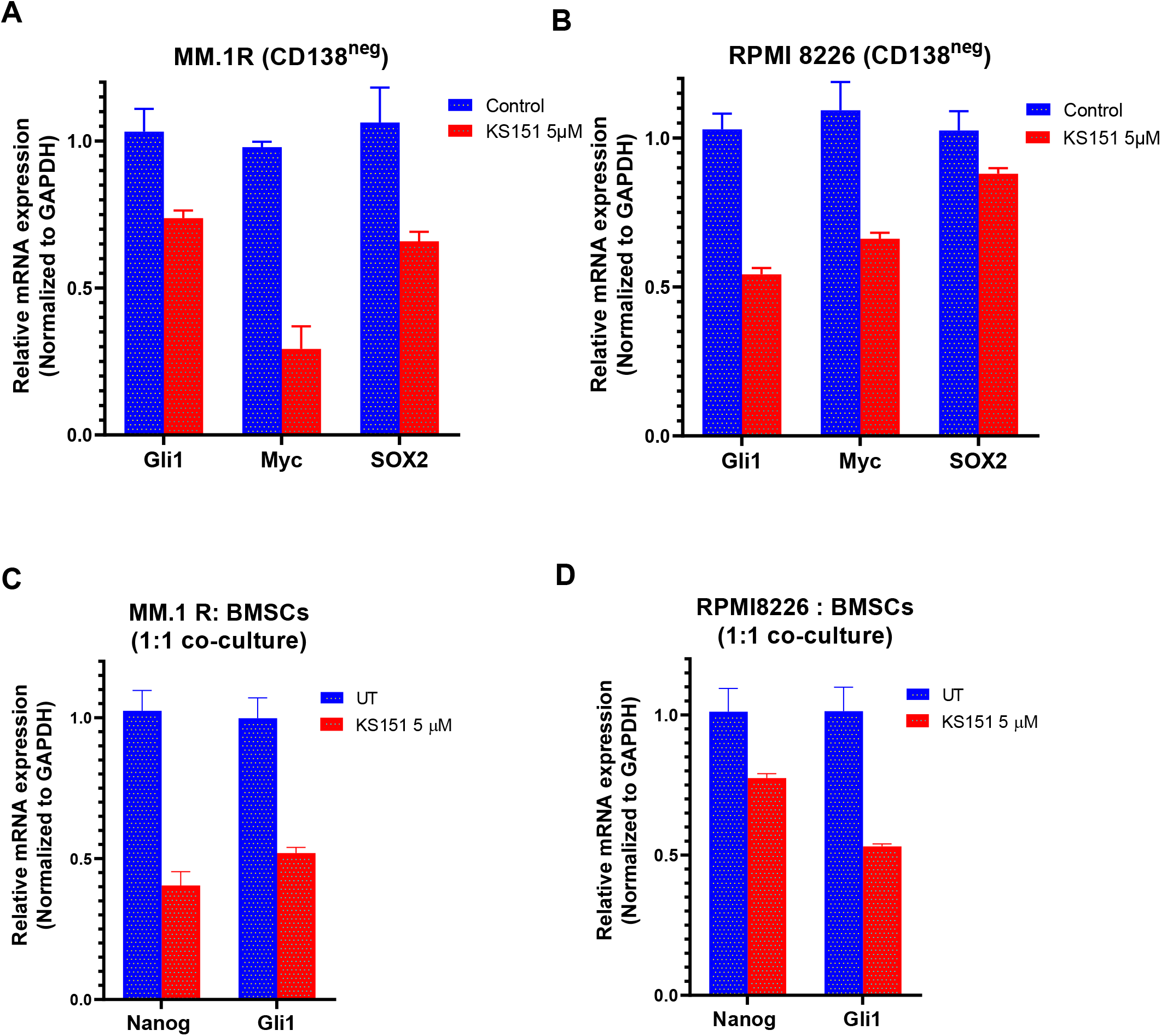
Effect of KS151 on stemness markers. ***A-B***, The effect of KS151 on CD138^neg^ stemness markers. MM.1R CD138^neg^ and RPMI 8226 CD138^neg^ cells were isolated using magnetic beads and treated with 5 μM KS151 for 24h then RNA was isolated, and RT-qPCR was performed using mentioned primers. ***C-D***, The effect of KS151 on stemness markers when MM.1R or RPMI 8226 were co-cultured with BMSCs. We plated 2 X10^5^ stromal cells in 6 well-plate for overnight, then 2 X10^5^ MM.1R or RPMI 8226 were added to them and treated with 5 μM of KS151 for 24h. RNA was isolated and RT reaction was followed by a qPCR.

## DISCUSSION

Despite the recent development of new treatment options, MM remains uniformly fatal because of intrinsic or acquired drug resistance [4, 65–68]. The key player in MM resistance to treatment is a subgroup of MM cells with stem cell-like properties called multiple myeloma stem cell-like cells [5]; therefore, it is reasonable to consider that any effective treatment for MM will have to be effective at targeting and eliminating the pool of MM stem cell-like cells. It has been demonstrated that stem cell-like cells overexpress BTK, and correlated to acquired drug resistance [17]. Moreover, BTK contributes to multidrug resistance [69]. Ibrutinib, a first-in-class, once-daily oral inhibitor of BTK, is approved by the FDA for the treatment of B cell malignancies [35, 36]. Ibrutinib has been shown to synergize in killing malignant cells when combined with bortezomib and lenalidomide in MM patient cells [70]. Though patient’s response of ibrutinib is encouraging, nonetheless patients develop resistance because of acquired mutations, which often involve the mutations of Cys481 to Ser (C481S) in the kinase domain [41–43, 71]. The C481S mutation disrupts the covalent binding between BTK and ibrutinib, which leads to a loss of inhibition of BTK enzymatic activity that ultimately results in ibrutinib resistance [41–43, 45]. To overcome this barrier, we designed and synthesized a novel BTK specific inhibitor, KS151 that does not interact with C481 in the kinase domain of BTK as determined by *in silico* studies. We first compared the inhibition of kinase ability of KS151 to ibrutinib. KS151 showed higher kinase inhibition ability compared with ibrutinib. Then, we next examined the specificity of KS151 toward BTK by using a sandwich-based array for detecting 71 tyrosine kinase phosphorylation signatures. Among different kinases, we found that KS151 selectively inhibited BTK.

BTK phosphorylates PLCγ2, and induces Ca2+ mobilization [72]. Our data demonstrates that KS151 inhibits the phosphorylation of PLCγ2 suggesting that KS151 regulates BTK signaling pathways. Using confocal microscopy, we further tested the effect of KS151 on the phospho-BTK protein. Importantly, KS151 inhibits the nuclear translocation of phospho-BTK. Studies have demonstrated that spatio-temporal dynamic of BTK regulates B cell receptor signaling [73]. The mechanism by which BTK shuttles between the nucleus and the cytoplasm remain elusive, along these lines, Mohamed and Vargas *et al.* (2000) showed that the SH3 domain plays an important role in retaining BTK in the cytoplasm as its deleted mutant showed predominant nuclear localization of BTK [62]. Moreover, they found that shuttling of BTK between the nucleus and the cytoplasm was independent of tyrosine phosphorylation status [62]. Furthermore, Gustafsson *et al.* (2012) have identified ankryin repeat domain 54 protein (ANKRD54), also known as Liar, as a BTK nuclear export protein [74]. Mohammad et al. (2013) identified a negative BTK regulator, 14-3-3 ζ, which binds to phosphorylated serine or threonine residues (S51/T495) of BTK, to target it for ubiquitination and degradation. Ibrutinib inhibits the 14-3-3 ζ binding to BTK [73]. Our data demonstrates that KS151 inhibits the cytoplasmic expression of phospho-BTK, however further studies required to understand the mechanism.

We assessed the cytotoxic potential of KS151 on various MM cell lines. We found that KS151 was effective in a number of MM cell lines. Then, we compared the effect of this novel molecule to other drugs that are already approved and being in clinic. We found that KS151 is more effective than most of chemotherapeutic agents. Moreover, when we compared its effect with other BTK inhibitors on bortezomib resistant cells, we found that a lower dose of KS151 is needed for bortezomib-resistant cell killing compared with other BTK inhibitors. We evaluated BTK inhibition with KS151 in combination with some chemotherapeutic agents, such as bortezomib, lenalidomide and panobinostat as a therapeutic strategy for chemo resistant MM stem cell-like cells (CD138^neg^). We demonstrated that KS151 augments the cytotoxic potential of therapeutic agents especially in MM stem cell-like cells. Similarly, combining lenalidomide with KS151 increased apoptosis of total MM.1R, and RPMI 8226, as well as CD138^neg^ RPMI 8226 and CD138^neg^ MM.1R cells. Moreover, combination of KS151 with panobinostat showed a better effect with higher doses of KS151 in MM.1R, RPMI 8226, CD138^neg^ MM.1R, and CD138^neg^ RPMI 8226.

Because BTK regulates the expression of stemness markers, we asked whether KS151 can inhibit the expression of stemness markers [17, 18]. KS151 was able to inhibit stemness markers in CD138^neg^ MM1.R and CD138^neg^ RPMI 8226 cells. Along these lines, a study demonstrates that knock down of BTK by shRNA inhibited the expression of Nanog in stem cell-like cells [17]. BMSCs secrete soluble factors that enhance the survival of MM [75], furthermore co-culturing BMSCs with MM cells increases resistance against chemotherapeutic agents and enhances the expression of stemness genes such as Nanog, SOX2, and OCT4 [18, 76]. Therefore, we investigated whether KS151 can modulate the expression of stemness markers when co-cultured with BMSCs. KS151 effectively inhibited the expression of Nanog and GLi1 even in the co-culture.

Overall, this study demonstrates that KS151 is a specific inhibitor of BTK that is more potent than ibrutinib and other chemotherapeutic agents, effective against bortezomib resistant cells, and potentiates the response of bortezomib, lenalidomide, and panobinostat in stem cell-like cells and regulates the expression of stem genes. Further experiments using animal models are needed to fully elucidate the mechanism of this novel agent.

## Acknowledgement

WOE designed, performed experiments, analyzed the data, and wrote the manuscript; OSO, and RJ performed experiments and analyzed the data. JKS performed docking studies; SKS, SCJ, SA, GPR and KG conceived ideas, contributed to providing essential reagents and synthesis of the analogs, and critically evaluating the study; TBA and MKP conceived ideas, designed the study, interpreted the results, and wrote the manuscript. JKS and SKS acknowledge the support of Multiscale Simulation Research Center (MSRC), Manipal University Jaipur for computational work. The synthesis of the analogs was partly supported by National Institutes of Health Grant ES028244, sub-awarded to SA and KG. This work was supported by the Camden Research Initiative Fund (MKP, TBA, and SJ), and an interdepartmental fund from Cooper Medical School of Rowan University, Camden, NJ (MKP). The authors are grateful to Marina N. Farag for composition of figures and providing technical assistance. The authors also thank Rachel King, Cooper Medical School of Rowan University Library, for careful proofreading and editing of the manuscript.

